# Can you make morphometrics work when you know the right answer? Pick and mix approaches for apple identification

**DOI:** 10.1101/288175

**Authors:** Maria D. Christodoulou, Nicholas H. Battey, Alastair Culham

## Abstract

Morphological classification of living things has challenged science for several centuries and has led to a wide range of objective morphometric approaches in data gathering and analysis. In this paper we explore those methods using apple cultivars, a model biological system in which discrete groups are pre-defined but in which there is a high level of overall morphological similarity. The effectiveness of morphometric techniques in discovering the groups is evaluated using statistical learning tools. No one technique proved optimal in classification on every occasion, linear morphometric techniques slightly out-performing geometric (72.6% accuracy on test set versus 66.7%). The combined use of these techniques with post-hoc knowledge of their individual successes with particular cultivars achieves a notably higher classification accuracy (77.8%). From this we conclude that even with pre-determined discrete categories, a range of approaches is needed where those categories are intrinsically similar to each other, and we raise the question of whether in studies where potentially continuous natural variation is being categorised the level of match between categories is routinely set too high.

## Introduction

From hominid stone implement design [1] to the identification of fossil sharks from their teeth [2], the extensive development of morphometric tools in the past few decades [3–5], has resulted in many exciting discoveries across scientific disciplines. In areas such as forensics and palaeontology, morphometrics may be the only tool available to researchers [2,6]. For over 2000 years morphology has remained the primary tool for field classification [7] although the tools used to gather data and analyse them have changed substantially. The classification of objects in general is a natural reaction of humans to the complexity of the world that surrounds them. Humans excel at pattern matching [8], a skill often exploited for security systems [9,10] and essential to classification. Arguably, this tendency can result in pareidolia, the misclassification of features to fit a preconceived model of limited scope [11]. Nevertheless, advanced pattern matching remains a crucial tool for navigating day to day life [12]. Many of the uses of pattern recognition (e.g. number plate reading [13] in carparks) rely heavily on statistical classification techniques, and have many potential biological applications [14–19]. Here we explore non-destructive morphometric sampling for classification of apple cultivars, the identity of which traditionally relies on the expert knowledge of a very small number of highly trained individuals yet the correct classification of an apple has immediate economic impact.

Continuous development in collection, recording, and analysis methods has given morphology a very sophisticated toolkit for taxonomists. Many recent taxonomic publications have exploited morphology under the umbrella of integrative taxonomy [20] which relies on the use of multiple data sources for inference [20]. The techniques most commonly combined with morphometrics are molecular [21], but can also include cytometry [20,22], chromosome counts [22] or the chemical composition of secreted compounds [23], all of which involve destructive sampling. The most important aspect of integrative taxonomy is the use of the appropriate data sources for the organisms in question. Combination of morphometrics and molecular markers can prove very successful in the delimitation of closely related taxa, both within botanical [24] and zoological [25–27] research. This success is taxon and technique dependent, as illustrated by the absence of morphometric resolving power in the works by Mamos et al. [28] and Lecocq et al. [23]. Diagnostic characters are often difficult to determine and quantify, and the selection process is challenging. Some examples of this difficulty include selecting the appropriate life stage[29]-contrasting larval stages to adults on *Culex* species - or morphological character - contrasting overall shape to specific landmarks on *Cobitis* populations [30]. Although the majority of these examples focus on shape description and quantification, colour may also be a vital source of morphometric data [21,26].

With more than 7,000 apple cultivars described [31](some authors estimate 10,000 cultivars [32]), fruit of all shapes, sizes, colours, flavour, and texture exist. This diversity makes identification a challenging task. Talented human experts can take years to master cultivar identification. By studying both internal and external morphological characters, apple experts rely on their in-depth knowledge of hundreds of cultivars, contextual awareness, and their understanding of biological variation within those cultivars to classify unknown samples [33–35]. They also commonly analyse their observations in a flexible manner, focusing on some aspects of the morphology more heavily in some cases than in others. To illustrate this, we present the hypothetical case of an expert identifying an apple that is uniformly dark red. In that case the expert would not consider cultivars which are almost exclusively green or yellow in colour, such as ‘Granny Smith’ and ‘Golden Delicious’, even if the shape and size of the sample fruit matches those cultivars; the expert would simply ignore the similarities in shape and focus on shape characters for apples that can be dark red in colour.

The fundamental challenge in identification of an individual apple by an expert is much greater than that, for instance, of identification of many bird species which can be done routinely at great distance using binoculars, due to the presence of consistent landmarks of shape, size, colour, etc. Fine-grained recognition algorithms are successful in identifying different species of birds in a variety of environments and from a variety of angles because the object being identified is fundamentally consistent in size, shape, and colour [36]. Similarly, the consistency of size, shape, and colour in flowers of the same taxon leads to routine benchmarking of fine-grained algorithms against floral datasets [36,37]. This has caused these characters to be used in extensively plant classification. Even the very well-studied British flora, has only recently gained an identification guide that does not depend on features flowers provide [38], despite the fact that experienced field botanists have long been able to identify plants in a vegetative state through knowledge and intuition. In the case of apples there is a need to identify individual fruit separated from the parent tree. The identification is at the level of cultivar and not species, and therefore the expected level of difference is small. As such it becomes crucial to standardise the imaging approach of the apples, such that variation detected is that of the fruit and not of its surroundings and the angle at which it is viewed.

Apple variety identification provides an ideal model to test the limits of morphological classification in biology because apple cultivars are usually clones and therefore the variation found is likely to be environmental in cause, and not genetic. By analysing clonal cultivars, we can be confident that there is a single correct answer to any identification. Both the challenge and novelty of this work is to discover whether apple cultivars can be identified accurately and reliably based on visual cues alone, in the absence of taste and smell. The challenge closest to our work is the collection of studies by Corney and colleagues [39–41] on automatic classification tools for *Tilia* leaves. The absence of sufficient landmarks for apple cultivars inspired us to study them from first principles, returning to basic morphometric tools and concepts in order to design a classification protocol.

Here we aim to discover whether the currently available arsenal of morphometric approaches is capable of grouping individual apples into their correct cultivar. We demonstrate that through the use of combined approaches a success rate of 78% can be achieved in this particularly challenging biological identification problem.

## Materials and methods

Fruit of twenty-seven apple cultivars were collected at the National Fruit Collection in Brogdale, Kent during the 2013 and 2014 growing seasons. These were collected when considered ready to harvest by the professional pickers, who routinely use appearance and flavour as indicators of ripeness. The list of cultivars sampled is presented in Table S1. Maximum length and maximum diameter were measured for each fruit using Vernier callipers (Mitutoyo Corporation, Japan). Weight, after removal of pedicel, was measured using precision scales calibrated to 0.01g (Denver Instrument S-402, New York). All measurements were made within 24 hours of harvest.

Each apple was placed against a blue (RGB: 0, 0, 255) background on a Kaiser Phototechnik R1 photographic stand and was photographed using a Nikon D5100 camera with a Nikon AF-S 40mm Micro NIKKOR f/2.8 DX G lens. The blue background was selected because it would interact to the smallest degree with apple skin colour, which is predominantly a combination of red and green pixels. The camera was positioned 0.50 m above the base of the stand, a setting that was not altered during the data collection and allowed capture of the entire outline of even the largest apples in the sample, at the same time retaining sufficient resolution for detailed digitisation. Each fruit was photographed a total of six times: one image for the calyx end, one for the pedicel end and four side-images (fruit rotated by 90° clockwise for every image), resulting in a total of 3,240 images (original image dimensions 4928×3264 pixels). Of the four side-images per fruit only the first two (the original and the 90° rotation from the original) were unique in terms of shape, the other two being their mirror images.

On each image, landmarks were recorded manually using the tpsDig2 software [42]. Landmark selection relied on the ability to consistently obtain the same landmarks on all the fruit. By observing collections of images from each cultivar six landmarks were selected for the digitisation: two on the crown apices, two on the shoulder apices, one on the calyx and one on the pedicel attachment point (illustrated in Figure 1).

**Fig 1:**
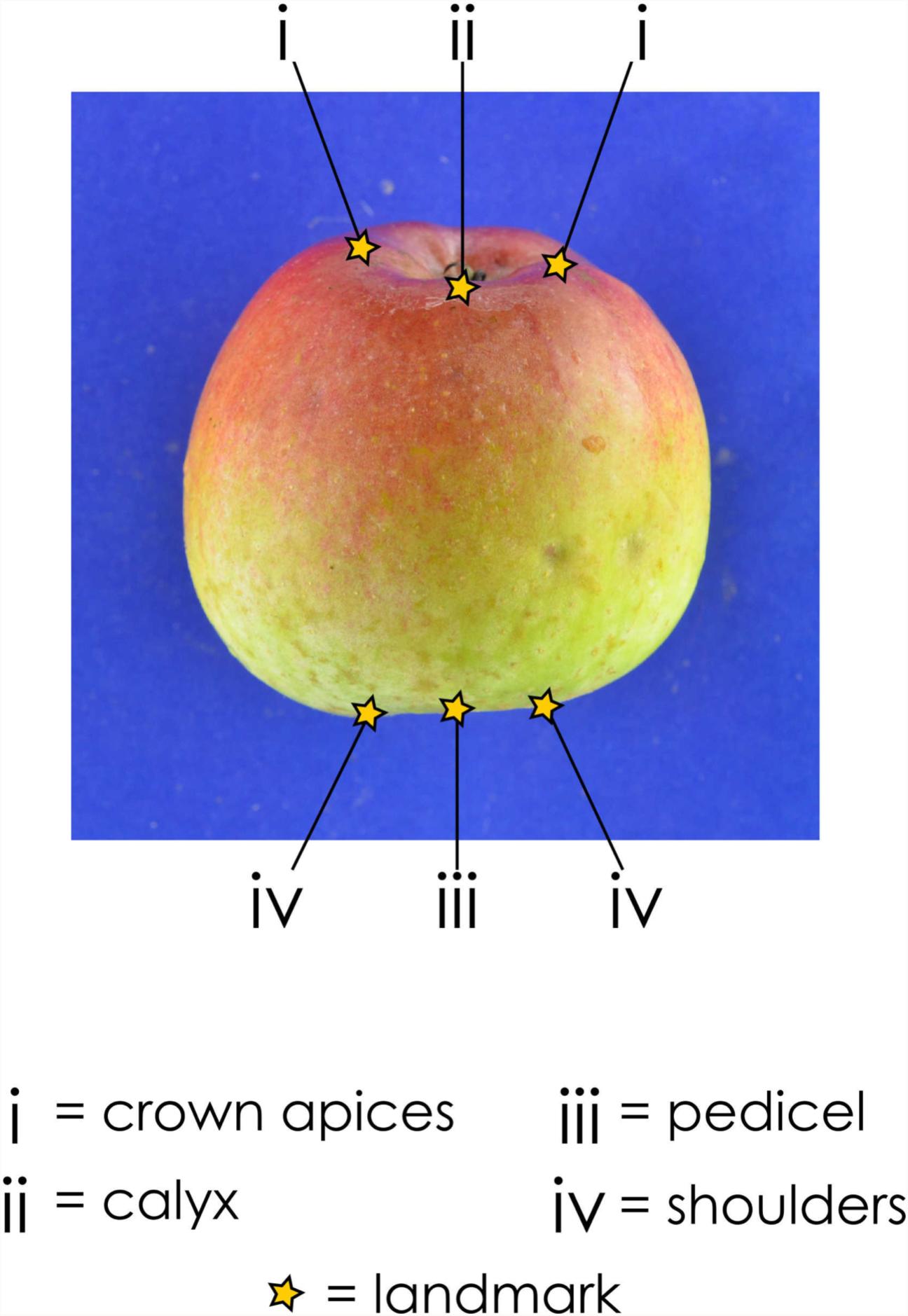
Selected landmarks for the geometric morphometrics dataset. Six landmarks were selected per image: two on the crown apices, one on the calyx, one on the pedicel attachment point and two on the shoulder apices.

To establish the degree of digitisation error, all 3,240 images were digitised twice, with a two-week gap, to ensure that the second digitisation was not affected by muscle memory. Analysis of digitisations was conducted using MorphoJ [43]. Digitisation error was calculated using Procrustes ANOVAs and found to be negligible across all samples. The first image of each fruit could have been used exclusively to describe its shape. This, however, would ignore the variation that the 90° rotation could provide. To be able to include the variation from the two views as well as to standardise between fruit, the landmark positions from the two views after a Procrustes superimposition were averaged. This process was repeated for the 180° and 270° views and the two datasets were then compared to establish possible digitisation error, which was also found to be negligible. After the Procrustes superimposition, the centroid size for each fruit was recorded. The Procrustes coordinates were then used to perform a Principal Components Analysis (PCA), the scores from which were recorded for each fruit.

Colour measurements were obtained by estimating the overall Red, Green and Blue (RGB) intensities per pixel for each image using ImageJ [44]. To reduce the dimensionality of the RGB colour measurements and remove the variation caused by the auto-white balance, a PCA was performed and the first principal component was retained as the overall colour measurement. The calyx images for each fruit were used to measure calyx area and the calyx “eye” (an opening in the calyx). This was performed using the tpsDig2 [42] by manually outlining the relevant edges.

From these measurements, two datasets were compiled, a linear morphometrics dataset including: maximum length; maximum diameter; weight; first principal component of colour; calyx area, and calyx (“eye”) aperture area, and a geometric morphometrics dataset including: weight; first principal component of colour; calyx area; calyx (“eye”) aperture area; Principal Component scores of Procrustes Coordinates, and centroid size. The datasets were then separated into training and testing sets with a 75-25% [45] allocation respectively using identical partitions for comparability. The training sets were then used on 12 classifiers (Table 1). Using the same partition for both datasets ensured that the accuracy estimate for the test set of the best performing linear morphometrics classification was directly comparable to the accuracy estimate for the test set of the best performing geometric morphometrics classification.

**Table 1:** Classifiers used to analyse linear and geometric morphometric datasets.

| Classifier | Abbreviation |
| --- | --- |
| Adaptive Mixture Discriminant Analysis [46] | AMD |
| Bagged Classification and Regression Tree [47] | BCART |
| C5.0 Classification Tree [48] | C5.0 |
| Classification and Regression Tree [49] | CART |
| Conditional Inference Random Forest [50] | CIRF |
| Feature Selection Random Forest [51] | FSRF |
| K-nearest Neighbor [52] | KNN |
| Naïve Bayes [53] | NB |
| Neural Network [54] | NN |
| Penalized Discriminant Analysis [55] | PDA |
| Robust Discriminant Analysis [56] | RDA |
| Random Ferns [57] | RF |
| Support vector Machine [58] | SVM |

Training used three repeats of 10-fold cross-validation. Each classifier was then tested using the test set and the classification confusion matrix, as well as the accuracy, kappa value, positive predictive rate, negative predictive rate, specificity, and sensitivity values were recorded. Paired t-tests on accuracy and kappa values were performed to compare between classifiers. The final model for each classification technique was selected in terms of highest accuracy and kappa values. All classification analysis was performed using the caret (Classification and Regression Training) package [59] in R [60].

To emulate the flexibility in character weighting shown by experts, who for instance might swap between using colour and size as a primary classifier, an ensemble approach was taken. When different datasets were used to train multiple classifiers, the success of each classifier with each cultivar could be recorded. For an unknown fruit tested against all the trained classifiers, the reliability of each prediction was assessed based on the accuracy of each classifier for the predicted cultivar. This process is illustrated in Figure 2. This replicated part of the expert flexibility by permitting the use of different characters for each classifier.

**Figure 2:**
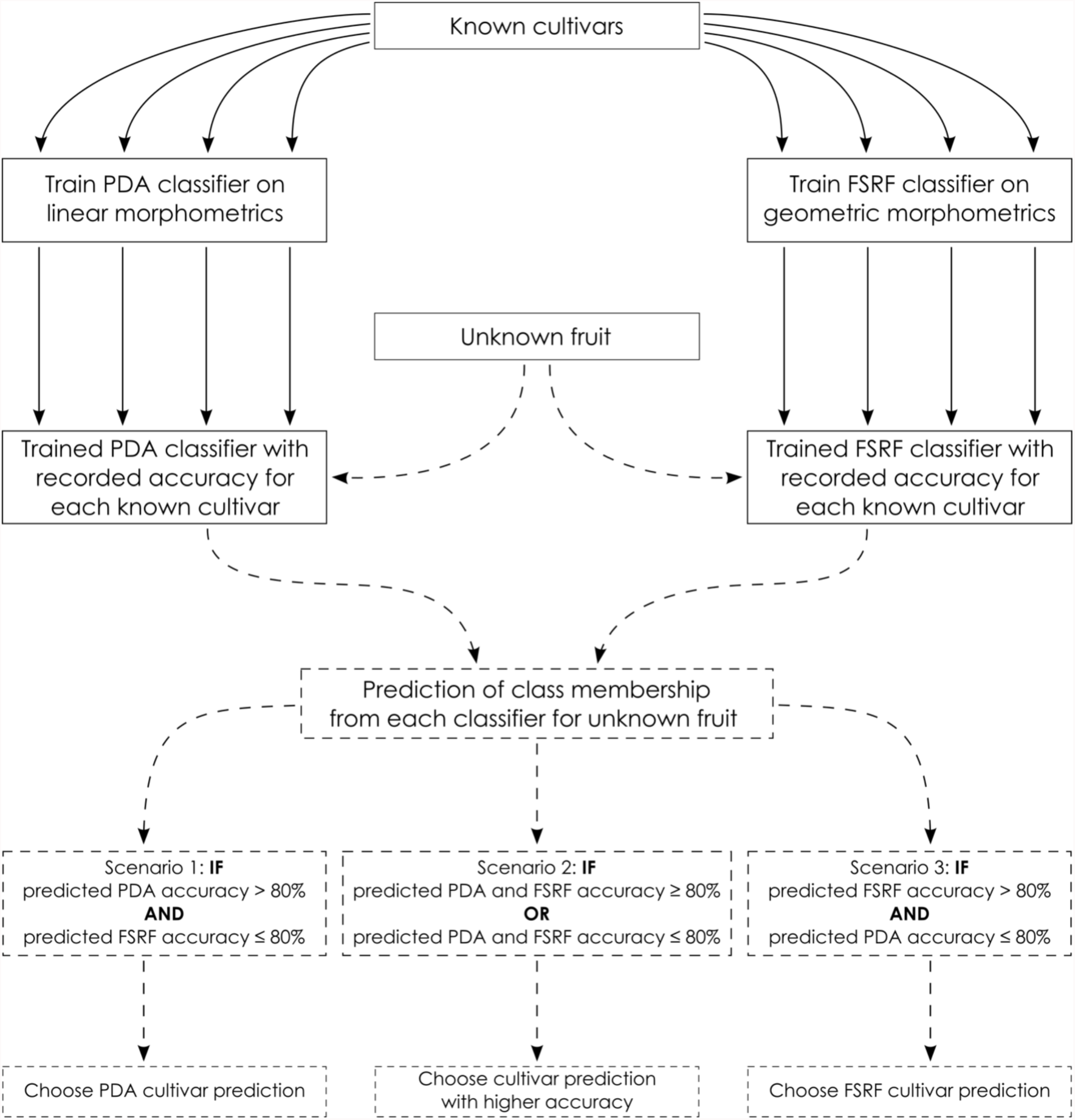
Flowchart of the manual ensemble process described in this work. After training the two classifiers and recording the cross-validation accuracy values for each cultivar, the unknown fruit is classified. The predicted class for the unknown fruit for each classifier is compared to the cross-validation accuracy for this class and the final prediction is selected according to which scenario applies.

As an alternative approach to the manual ensemble procedure, the linear and geometric morphometrics datasets were combined to create a “kitchen sink” [61] dataset, to investigate whether the concatenation of raw data led to a more successful classification. The concatenated dataset was partitioned in the same way as the linear and geometric morphometrics datasets.

All the images used in the above study are deposited in the Reading Apple Image Library, accessible through the University of Reading Herbarium webpages. Together with the fully matured fruit presented in this study, the Image library also contains standardised images for fruit sampled from 12 of the cultivars at different time points from anthesis (weekly for the first two months from anthesis, and fortnightly later on). For each time-point, ten fruit were sampled and photographed as described here. Additionally, longitudinal sections from calyx to stem were performed and each side was photographed. For six cultivars, sampling was repeated for a second year. This resulted in 13360 images for 27 cultivars.

## Results

Prior to training and testing of classifiers, the RGB colour values were reduced using principal component analysis (PCA). As the first principal component explained 94.3% of the overall colour variation, it was deemed a sufficient colour proxy, and was the only component retained for the remainder of the analysis. Classifier comparison was performed using accuracy and kappa values over the collected datasets. Classifiers were tested over four different settings: first against the linear morphometrics dataset, second against the geometric morphometrics one, third using the manual ensemble approach, and finally against the kitchen-sink dataset. The remainder of this section is split to accommodate these four approaches.

### Linear morphometrics

Of the 12 classifiers studied, Penalised Discriminant Analysis (PDA) had the highest mean accuracy and kappa values (accuracy: 73.0%, kappa: 0.722) for cross-validation of the training set and for this reason it was selected as the most appropriate classification technique. Following this, the test set, which comprised 135 fruit (5 from each cultivar) was analysed using the trained PDA classifier resulting in a percentage accuracy estimate over all classes (overall accuracy percentage) of 72.6%. Individual misclassifications for each fruit in the test set are in Tables S4-S6 in **Supplementary Materials**.

### Geometric morphometrics

Of the 11 classifiers tested, the best performing was the Feature Selection Random Forest (FSRF) as it had the highest mean accuracy and mean kappa values (accuracy: 66.5%, kappa: 0.654). Individual misclassifications for each fruit in the test set are in Tables S7 and S8 in **Supplementary Materials**.

### Manual ensemble

For manual ensemble, the predictions for the test set of the PDA on linear morphometrics were combined with the predictions of the FSRF classifier of the geometric morphometrics by using the accuracy estimates of cross-validation for each class (the detailed manual ensemble protocol is described in Materials and Methods). The confusion matrix for the test set classification, which is the per-class performance of the trained classifier, is illustrated in Figure 3. Through the use of a heat-map, Figure 3 contrasts the actual class (cultivar) to which each fruit in the test set belonged (Reference) against what class it was predicted as (Prediction) by the trained classifier. Correct classifications are on the diagonal of the heat-map, with darker shades of blue illustrating greater success rates.

**Figure 3:**
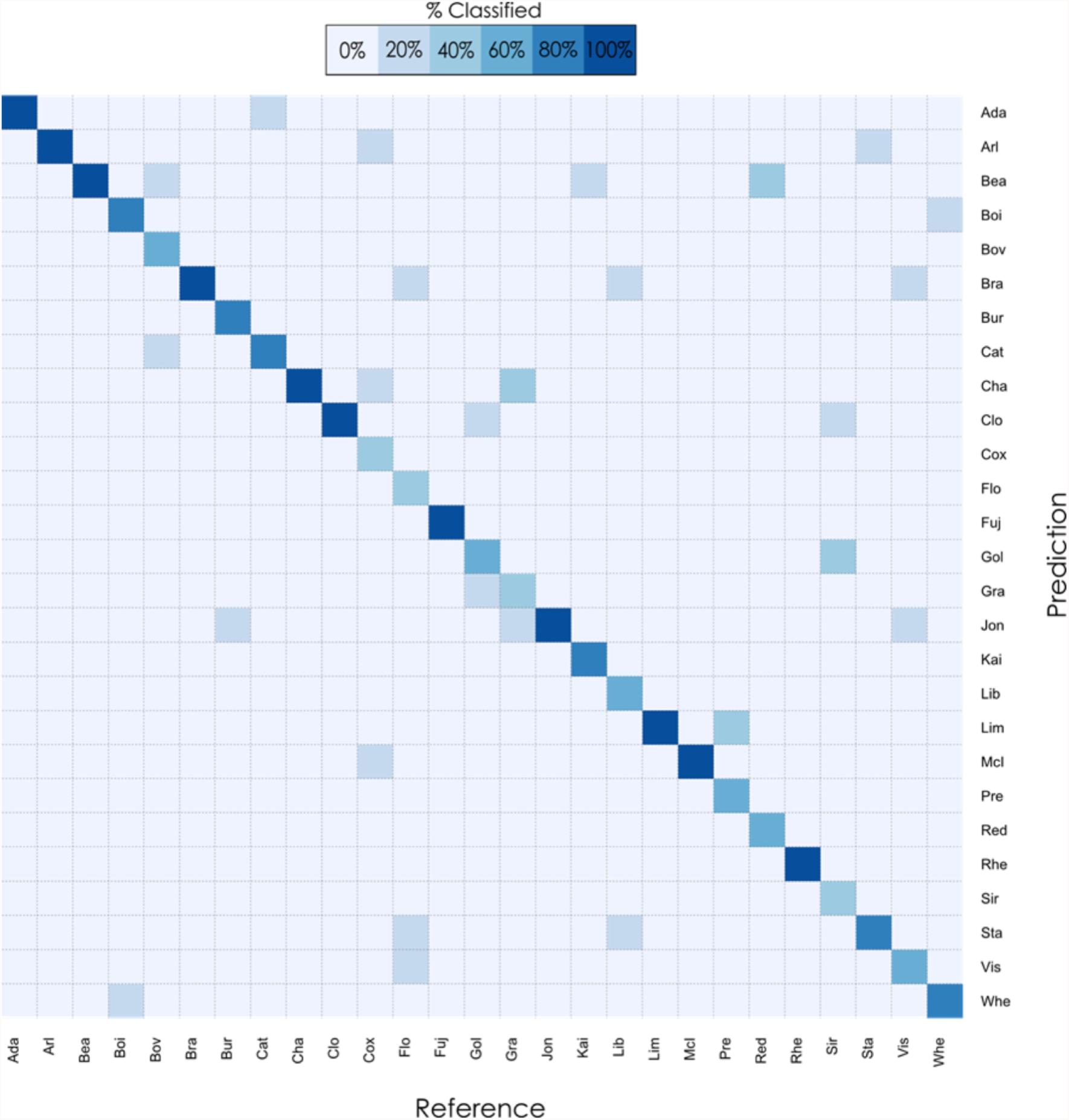
Confusion matrix from the Manual ensemble classification using the test set. The colours of the heat map correspond to the percentage of classification in each category. The accuracy obtained from the manual ensemble was 77.8% compared with 66.7% for the FSRF and 72.6% for the PDA on the same test set.

### Kitchen-sink

Of the 11 classifiers tested with the “kitchen sink” dataset, the best performing was the Adaptive Mixture Discriminant Analysis (AMD) with mean accuracy of 70.5% and a kappa value of 0.692 for cross-validation.

The predictions of the test set samples for each classifier by cultivar are summarised in Figure 4, which demonstrates that every cultivar could be correctly classified using one of the four techniques. If one technique failed to classify a cultivar, another often turned out to be successful. The success of the classification techniques varied between the cultivars. For example, in the case of ‘Adam’s Pearmain’ (Ada), all four approaches had a very high success rate, with three of them reaching 100% accuracy, and the lowest one reaching 80%. Findings were similar for ‘Cloden’ (Clo), with two methods reaching 100% accuracy and the remaining two 80%. Less successful was the case of ‘Bovarde’ (Bov), for which the FSRF classifier relying on the geometric morphometrics dataset failed to correctly identify any of the samples in the test set. Although none of the other classifiers succeeded in correctly identifying all five ‘Bovarde’ samples in the test set, they successfully identified three.

**Figure 4:**
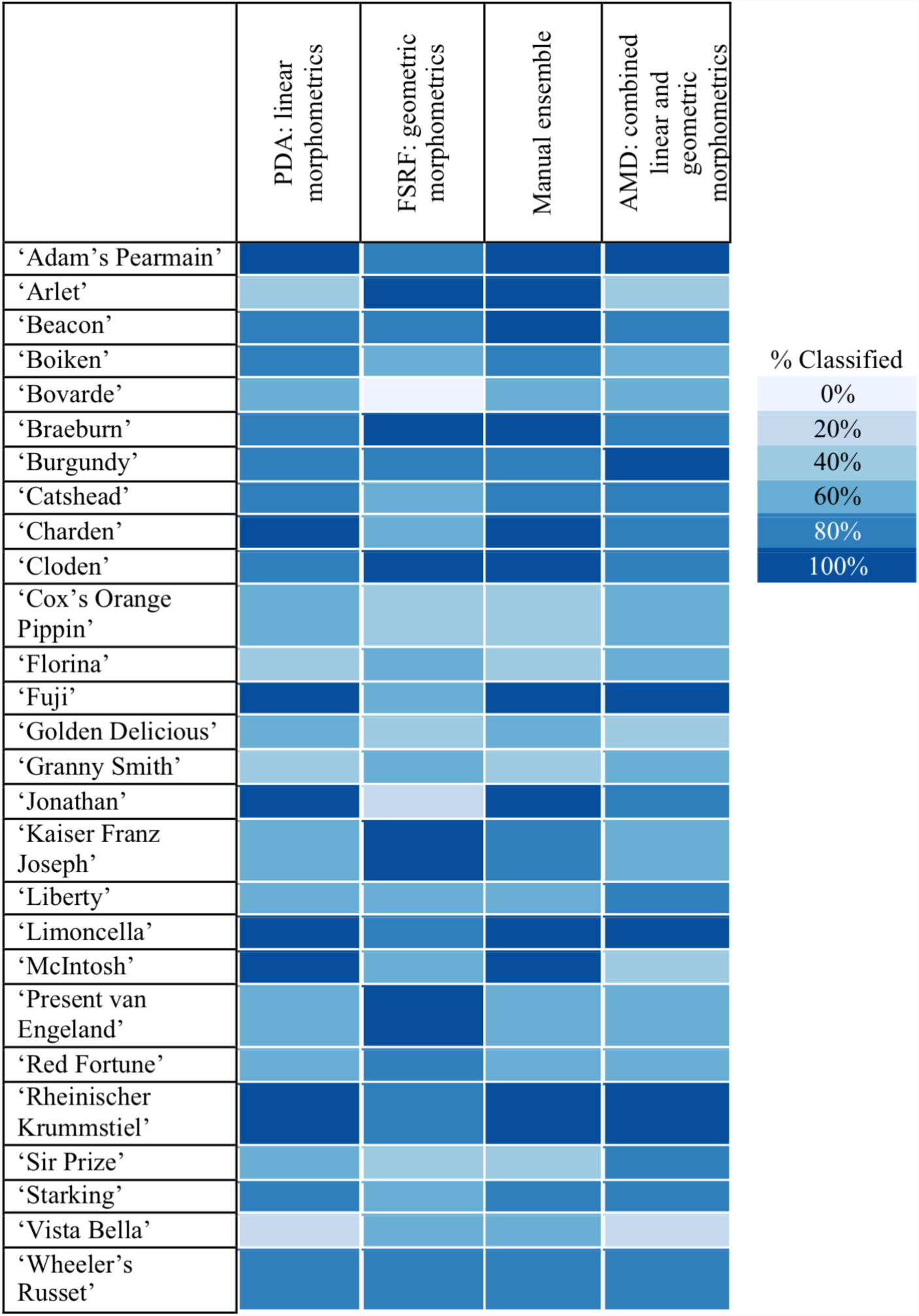
Summary of prediction rates of test set by cultivar for the classifiers using the same colours as the heatmap in Figure 3. Classifier abbreviations are explained in Materials and Methods.

## Discussion

The advantage of studying apple cultivars which are clonally propagated was that we could be certain of the correct classification for each individual apple. This contrasts with equivalent studies of variation in species because species are conceptual constructs which may change over time [62,63]. For instance Compton and Hedderson [64] required 17 morphometric variables to separate a single variable species into four distinct ones, and those supported by correlation with geographic distribution. Despite the clonal identity within apple cultivars and the variety of morphometric measurements used in this study, our classifications still resulted in misidentifications of many individual apples. Here we consider some of the underlying reasons for these.

We learned two major lessons during the process of automating classification.

### Lesson 1: There is no free lunch

The performance and choice of classifier depends on the nature of the underlying data. For example, using linear morphometric techniques the best performing classifier was a PDA (accuracy 72.6%); for geometric morphometrics it was a FSRF (accuracy 66.7%). This finding is consistent with the “No free lunch” theorem. Stated formally by Wolpert and Macready [65], the theorem suggests that the performance of all classifiers is equal when the totality of possible problems is considered. This means that for every classifier there exists a possible problem where that classifier outperforms every other classifier. In our study two different morphometric datasets created two different classification problems, each analysed most effectively by a different classifier. This strong interaction between dataset and classifier is one of many examples of the no-free lunch theorem. Adding to the complexity is the impact of cultivar as a variable on the classifier and dataset interaction. As demonstrated in Figure 4, some cultivars were more accurately identified using one classifier and others by another. This suggests that in addition to selecting the appropriate classifier for the dataset, it is important to establish for every cultivar how accurately each combination performs. To illustrate this, four apples (‘Arlet’, ‘Bovarde’, ‘Jonathan’, ‘Kaiser Franz Joseph’) which were all part of the test set, are shown in Figure 5.

**Figure 5:**
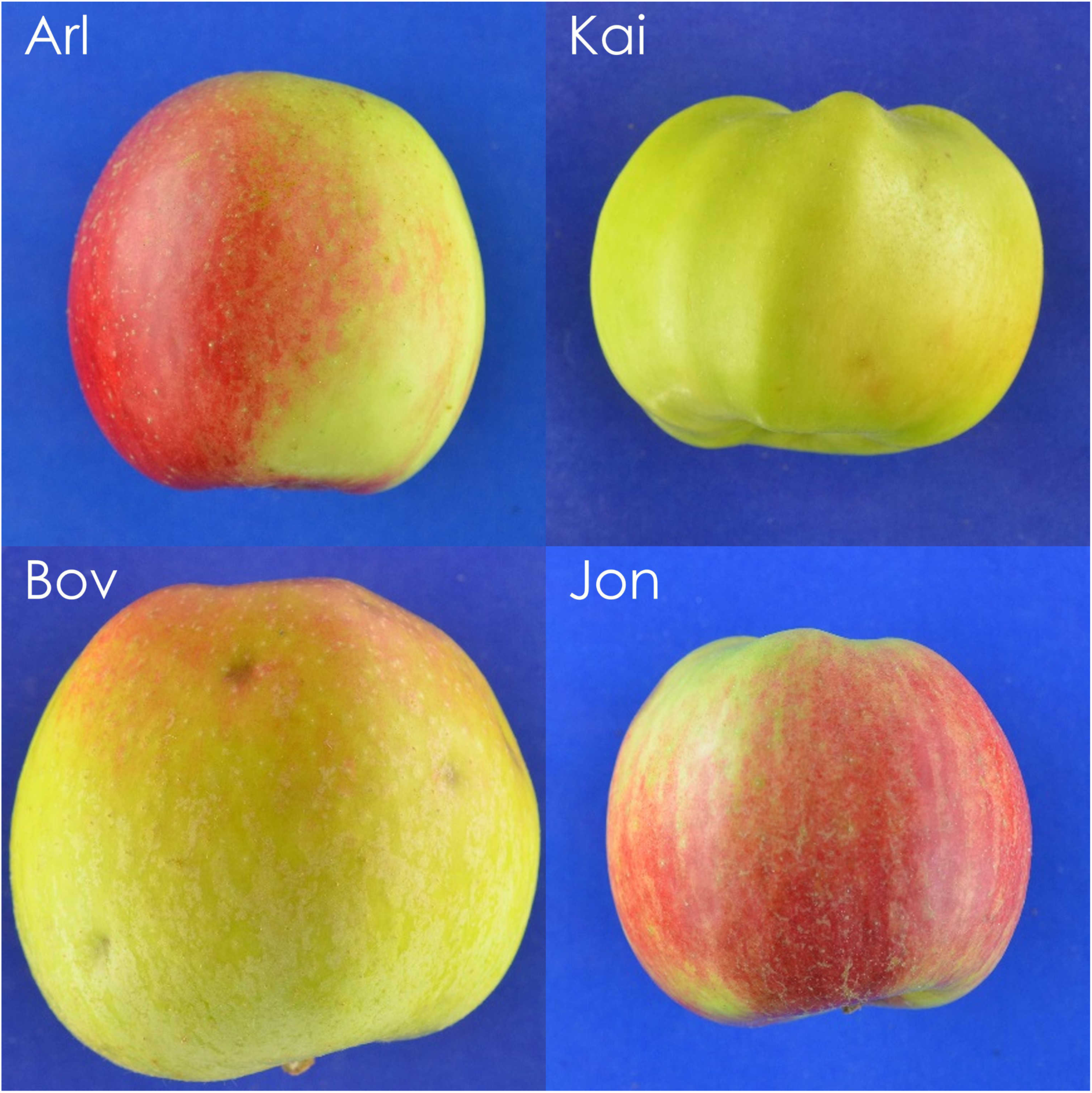
Four fruit examples that were misclassified by one of the two classifiers. In the top two rows Arl and Kai were misclassified by the PDA but were successfully classified by the FSRF. In the bottom two rows, Bov and Jon were misclassified by the FSRF but successfully classified by the PDA. The first image in each row is the misclassified apple, the second is the cultivar it was classified as, and the third one is an example of the training dataset for the correct classification.

All of ‘Arlet’ (Arl) and ‘Kaiser Franz Joseph’(Kai) samples in the test set were accurately classified using FSRF. Some ‘Arlet’ and ‘Kaiser Franz Joseph’ samples in Figure 5 were misclassified using PDA (which had 40% success rate for ‘Arlet’ and 60% for ‘Kaiser Franz Joseph’). ‘Jonathan’ (Jon) and ‘Bovarde’(Bov) were classified more accurately by the PDA than the FSRF (100% and 60% respectively with the PDA as opposed to 20% and 0% with FSRF). Why are some cultivars more identifiable using one classifier than with another? For the cultivars that performed better with geometric morphometrics, such as ‘Kaiser Franz Joseph’, we propose that the distinctive fruit geometry failed to translate into recorded parameters in linear morphometrics. For the cultivars that performed better with the linear morphometrics, such as ‘Jonathan’, we propose that the overall geometry of the fruit was not as distinctive as the length and diameter measurements.

### Lesson 2: Pick and mix

Improved accuracy results from the flexible combination of linear and geometric morphometrics classifiers. The successful protocol used as inspiration the flexibility of information that human identification experts can employ, by combining different data-sources (in this case linear and geometric morphometrics). The explanation for the superiority of this method (over both linear and geometric classifications) lies in the differences of accuracy per cultivar for each classification. The successful protocol gives different weights to the predictions depending on how accurate each classifier has been in the past for that particular prediction. For example, if an unknown fruit was predicted as ‘Jonathan’ by the FSRF and as ‘McIntosh’ by the PDA then the manual ensemble would classify it as a ‘McIntosh’ since the FSRF is weak at predicting ‘Jonathan’ (or ‘McIntosh’), whereas the PDA is strong for both cultivars. By using this method and effectively relying on each classifier for the cultivars they were good at, the classification performance improved to an overall 77.8%. As a technique, it was particularly effective when there was a marked difference in the classification accuracy for a cultivar (e.g. with ‘Jonathan’). When the classifiers performed at similar levels (e.g. ‘Florina’ with 40% with PDA and 60% with FSRF) then some accuracy could be lost in the classification (‘Florina’ in manual ensemble had 40% accuracy).

Although the “kitchen sink” approach was more accurate (71.1%) than the FSRF, it was less accurate than the PDA or manual ensemble. This indicates that the simple concatenation of both datasets increased noise. Aside from performance there was a fundamental difference between the “kitchen sink” and the manual ensemble. Both techniques used all the information available by including linear and geometric morphometrics but whereas the “kitchen sink” merged raw data, the manual ensemble exploited the strengths of each dataset.

## Conclusions

The primary objective of this work was to discover whether apple cultivars could be identified using automated processes by exploiting some of the strategies apple experts employ in combination with current morphometric approaches. We conclude that computers can effectively simulate the approach used by apple experts, prioritising some data over others, in a cultivar- and situation-specific way.

The most impactful novelty of this work is methodological; specifically, the use of explicit geometric and linear morphometrics in combination with statistical learning has great relevance to wider biological research in identification and classification. It is not clear why such an ensemble method is not routinely used for biological identification as it combines the strength of several approaches. Ensemble learning techniques started gaining popularity in the 1990s for statistical earning specifically because they can combine weak learners (classifiers with low accuracy) to create a strong learner (classifier with high accuracy) [54].

Modern plant taxonomy could embrace this approach and take advantage of current computing power. This would permit the re-evaluation of data-sources which on their own may only lead to weak learners, but in thoughtful combinations have the potential to provide novel insight into classification of the organism under study. Crucially, the incorporation of multiple datasets towards a single classification problem is not about simply combining raw data from multiple sources; it is about the careful integration of such data and multiple approaches to analysis to improve insight and understanding.

## Acknowledgments

We would like to thank Professor Richard Sibly, Dr Louise Johnson, Dr Tom Oliver, and Dr Jonathan Clark for constructive feedback on early drafts of this manuscript. We also thank Dr Matthew Ordidge for assistance with apple sampling, and Dr Kálmán Könyves for help with graphics.

