## Supplementary Materials for "Can you make morphometrics work when you know the right answer? Pick and mix approaches for apple identification"

Supplementary material

Table S1: Selected cultivars for morphological classification study.

| Cultivar | Abbreviation | Harvest Date |
|---|---|---|
| ‘Adam’s Pearmain’ | Ada | 23/09/2013 |
| ‘Arlet’ | Arl | 11/09/2014 |
| ‘Beacon’ | Bea | 02/09/2013 |
| ‘Boiken’ | Boi | 07/10/2013 |
| ‘Bovarde’ | Bov | 07/10/2013 |
| ‘Braeburn’ | Bra | 11/09/2014 |
| ‘Burgundy’ | Bur | 11/09/2014 |
| ‘Catshead’ | Cat | 23/09/2013 |
| ‘Charden’ | Cha | 11/09/2014 |
| ‘Cloden’ | Clo | 11/09/2014 |
| ‘Cox’s Orange Pippin’ | Cox | 11/09/2014 |
| ‘Florina’ | Flo | 11/09/2014 |
| ‘Fuji’ | Fuj | 07/10/2013 |
| ‘Golden Delicious’ | Gol | 11/09/2014 |
| ‘Granny Smith’ | Gra | 11/09/2014 |
| ‘Jonathan’ | Jon | 11/09/2014 |
| ‘Kaiser Franz Joseph’ | Kai | 23/09/2013 |
| ‘Liberty’ | Lib | 11/09/2014 |
| ‘Limoncella’ | Lim | 07/10/2013 |
| ‘McIntosh’ | McI | 11/09/2014 |
| ‘Present van Engeland’ | Pre | 23/09/2013 |
| ‘Red Fortune’ (Sport of ‘Fortune’) | Red | 02/09/2013 |
| ‘Rheinischer Krummstiel’ | Rhe | 07/10/2013 |
| ‘Sir Prize’ | Sir | 11/09/2014 |
| ‘Starking’ (Sport of ‘Delicious’) | Sta | 11/09/2014 |
| ‘Vista Bella’ | Vis | 17/07/2014 |
| ‘Wheeler’s Russet' | Whe | 07/10/2013 |

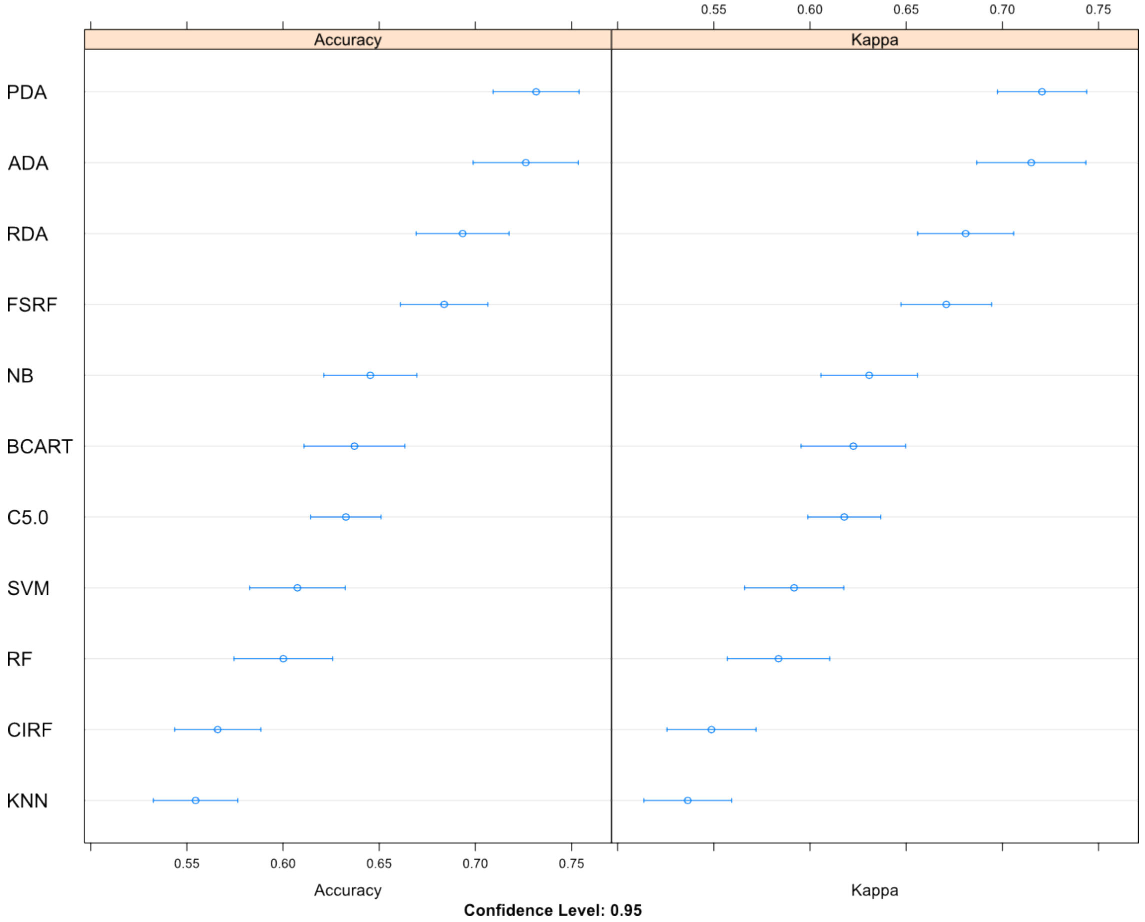


Figure S1: Accuracy and kappa value results with 95% confidence interval for the classification methods on the linear morphometrics dataset.

Table S2: Summary table of results from paired t-test comparisons between classifiers for linear morphometrics dataset. The results above the diagonal (represented by grey cells) were derived from t-tests on the accuracy values. The results below the diagonal were derived from t-tests on the kappa values. Significance is indicated with * levels (NS: Not Significant, *:p≤0.05, **≤0.01, ***≤0.001).

| | | t-tests on accuracy | | | | | | | | | | | |
|---|---|---|---|---|---|---|---|---|---|---|---|---|---|
| | | AMD | PDA | RDA | NB | C5.0 | FSRF | CIRF | RF | BCART | KNN | SVM | NN |
| t-tests on kappa | AMD | | NS | * | *** | *** | NS | *** | *** | *** | *** | *** | *** |
| | PDA | NS | | * | *** | *** | ** | *** | *** | *** | *** | *** | *** |
| | RDA | * | * | | NS | ** | NS | *** | *** | ** | *** | *** | *** |
| | NB | *** | *** | NS | | NS | NS | *** | NS | NS | *** | NS | *** |
| | C5.0 | *** | *** | ** | NS | | *** | ** | NS | NS | ** | NS | *** |
| | FSRF | NS | ** | NS | NS | *** | | *** | *** | ** | *** | *** | *** |
| | CIRF | *** | *** | *** | *** | ** | *** | | NS | ** | NS | NS | *** |
| | RF | *** | *** | *** | NS | NS | *** | NS | | NS | NS | NS | *** |
| | BCART | *** | *** | ** | NS | NS | ** | ** | NS | | * | NS | *** |
| | KNN | *** | *** | *** | *** | ** | *** | NS | NS | * | | NS | *** |
| | SVM | *** | *** | *** | NS | NS | *** | NS | NS | NS | NS | | *** |
| | NN | *** | *** | *** | *** | *** | *** | *** | *** | *** | *** | *** | |

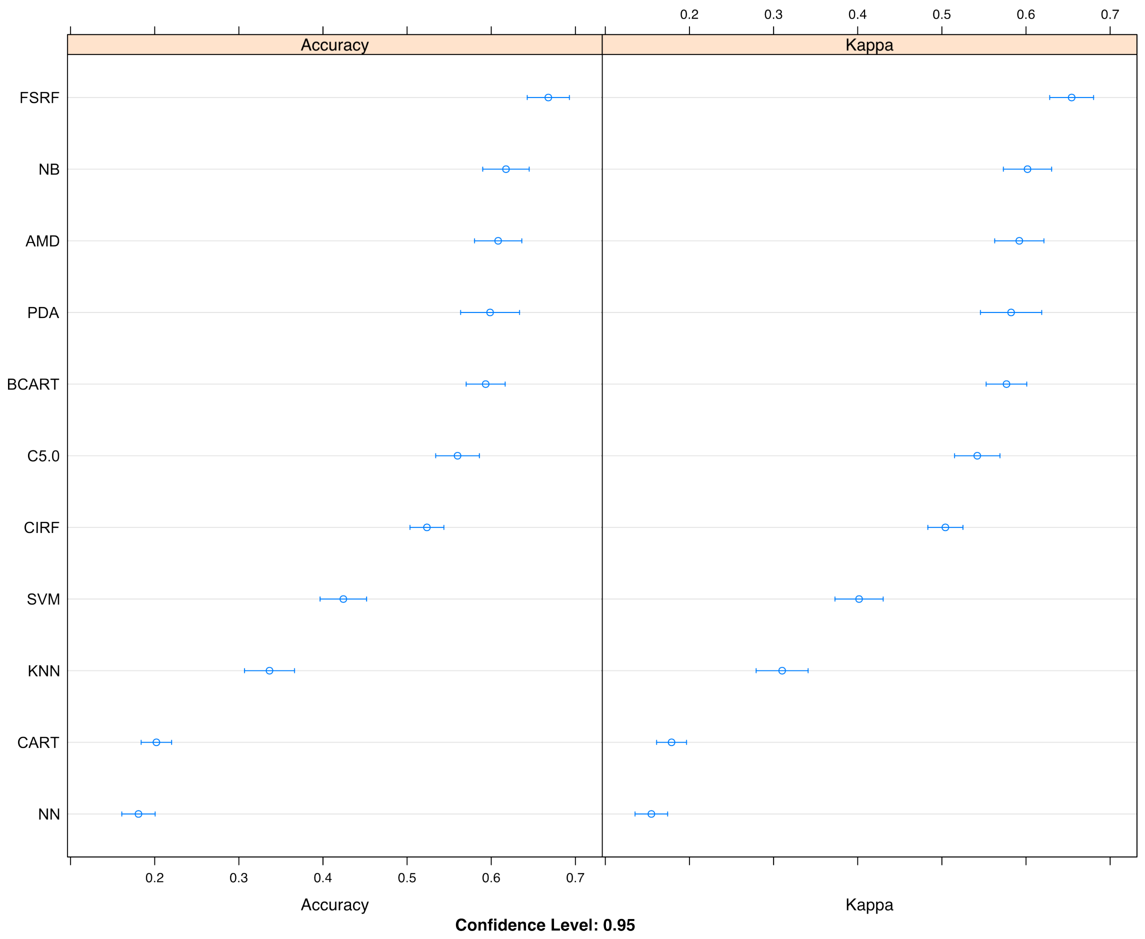


Figure S2: Accuracy and kappa value results with 95% confidence interval for the classification methods on the geometric morphometrics dataset.

Table S3: Summary table of results from paired t-test comparisons between classifiers for the geometric morphometrics dataset. The results above the diagonal (represented by grey cells) were derived from t-tests on the accuracy values. The results below the diagonal were derived from t-tests on the kappa values. Significance is indicated with * levels (NS: Not Significant, *:p≤0.05, **≤0.01, ***≤0.001).

| | | t-tests on accuracy | | | | | | | | | | |
|---|---|---|---|---|---|---|---|---|---|---|---|---|
| | | AMD | PDA | NB | FSRF | CIRF | C5.0 | BCART | KNN | SVM | NN | CART |
| t-tests on kappa | AMD | | NS | NS | NS | * | NS | NS | *** | *** | *** | *** |
| | PDA | NS | | NS | NS | * | NS | NS | *** | *** | *** | *** |
| | NB | NS | NS | | NS | *** | NS | NS | *** | *** | *** | *** |
| | FSRF | NS | NS | NS | | *** | *** | * | *** | *** | *** | *** |
| | CIRF | * | * | *** | *** | | NS | * | *** | *** | *** | *** |
| | C5.0 | NS | NS | NS | *** | NS | | NS | *** | *** | *** | *** |
| | BCART | NS | NS | NS | * | * | NS | | *** | *** | *** | *** |
| | KNN | *** | *** | *** | *** | *** | *** | *** | | *** | *** | *** |
| | SVM | *** | *** | *** | *** | *** | *** | *** | *** | | *** | *** |
| | NN | *** | *** | *** | *** | *** | *** | *** | *** | *** | | NS |
| | CART | *** | *** | *** | *** | *** | *** | *** | *** | *** | NS | |

Table S4: Percentage match for misclassifications (Part 1 of 3) from linear morphometrics using Penalised Discriminant Analysis. All cultivars with at least a single misclassification are included. The highest three percentages for each cultivar are included. If the correct classification is not in the top three then it is also included together with its rank. If the fruit was correctly classified only the correct classification posterior is included.

| Cultivar | Sample 1 | Sample 2 | Sample 3 | Sample 4 | Sample 5 |
|---|---|---|---|---|---|
| Arl | 1: Sta (30.95%)<br>2: Lib (21.74%)<br>3: Flo (20.41%)<br>8: Arl (4.13%) | 1: Sta (21.94%)<br>2: Lib (21.85%)<br>3 Flo (16.71%)<br>6: Arl (9.55%) | 1: Lib (41.59%)<br>2: Arl (27.18 %) | 1: Arl (34.55%) | 1: Arl (52.90%) |
| Boi | 1: Whe (99.18%)<br>2: Boi (0.59%) | 1: Boi (99.33%) | 1: Boi (82.91%) | 1: Boi (72.25%) | 1: Boi (98.80%) |
| Bov | 1: Red (45.23%)<br>2: Lim (32.66%)<br>3: Kai (21.53%)<br>8: Bov (0.01%) | 1: Cat (95.73%)<br>2: Boi (3.33%)<br>3: Bov (0.92%) | 1: Bov (76.29%) | 1: Bov (75.13%) | 1: Bov (57.43%) |
| Bur | 1: Jon (57.57%)<br>2: Lib (12.95%)<br>3: Bur (9.27%) | 1: Sta (48.76%)<br>2: Bur (24.78%) | 1: Bur (48.41%) | 1: Bur (64.94%) | 1: Bur (37.31%) |
| Cat | 1: Ada (98.62%)<br>2: Whe (0.97%)<br>3: Rhe (0.16%)<br>7: Cat (0.001%) | 1: Cat (94.21%) | 1: Cat (99.91%) | 1: Cat (99.99%) | 1: Cat (95.20%) |

Table S5: Percentage match for misclassifications (Part 2 of 3) from linear morphometrics using Penalised Discriminant Analysis. All cultivars with at least a single misclassification are included. The highest three percentages for each cultivar are included. If the correct classification is not in the top three then it is also included together with its rank. If the fruit was correctly classified only the correct classification posterior is included.

| Cultivar | Sample 1 | Sample 2 | Sample 3 | Sample 4 | Sample 5 |
| --- | --- | --- | --- | --- | --- |
| Clo | 1: Gol (54.94%)<br>2: Clo (31.85%) | 1: Clo (98.00%) | 1: Clo (97.76%) | 1: Clo (98.77%) | 1: Clo (61.24%) |
| Cox | 1: Arl (29.25%)<br>2: Jon (29.23%)<br>3: Bur (13.96%)<br>6: Cox (4.25%) | 1: Cox (68.81%) | 1: Cox (89.95%) | 1: Cox (36.49%) | 1: Cox (55.88%) |
| Flo | 1: Vis (20.01%)<br>2: Jon (17.44%)<br>3: Flo (15.45%) | 1: Bur (24.37%)<br>2: Vis (22.58%)<br>3: Flo (20.01%) | 1: Sta (32.51%)<br>2: Flo (28.56%) | 1: Flo (43.49%) | 1: Flo (80.46%) |
| Gol | 1: Clo (46.58%)<br>2: Gra (36.32%)<br>3: Gol (9.12%) | 1: Gra (65.49%)<br>2: Gol (20.94%) | 1: Gol (72.84%) | 1: Gol (55.49%) | 1: Gol (62.77%) |
| Gra | 1: Jon (32.32%)<br>2: Sta (27.92%)<br>3: Lib (2.09%)<br>11: Gra (0.01%) | 1: Cha (57.52%)<br>2: Clo (29.43%)<br>3: Gra (11.73%) | 1: Cha (57.92%)<br>2: Gra (29.64%) | 1: Gra (83.52%) | 1: Gra (80.63%) |

Table S6 Percentage match for misclassifications (Part 3 of 3) from linear morphometrics using Penalised Discriminant Analysis. All cultivars with at least a single misclassification are included. The highest three percentages for each cultivar are included. If the correct classification is not in the top three then it is also included together with its rank. If the fruit was correctly classified only the correct classification posterior is included.

| Cultivar | Sample 1 | Sample 2 | Sample 3 | Sample 4 | Sample 5 |
|---|---|---|---|---|---|
| Kai | 1: Red (95.56%)<br>2: Kai (2.35%) | 1: Red (70.94%)<br>2: Kai (23.98%) | 1: Kai (77.63%) | 1: Kai<br>(99.44%) | 1: Kai<br>(76.87%) |
| Lib | 1: Sta (34.28%)<br>2: Lib (26.65%) | 1: Bra (33.2%)<br>2: Lib (28.24%) | 1: Lib (53.28%) | 1: Lib<br>(51.97%) | 1: Lib<br>(64.80%) |
| Pre | 1: Lim (51.72%)<br>2: Pre (48.21%) | 1: Pre (99.99%) | 1: Pre (53.98%) | 1: Pre<br>(99.99%) | 1: Pre<br>(99.99%) |
| Sir | 1: Gol (45.35%)<br>2: Gra (27.82%)<br>3: Cox (13.58%)<br>7: Sir (1.09%) | 1: Gol (38.08%)<br>2: Gra (35.65%)<br>3: Sir (19.11%) | 1: Sir (96.34%) | 1: Sir<br>(96.25%) | 1: Sir<br>(82.12%) |
| Sta | 1: Arl (74.73%)<br>2: Lib (13.47%)<br>3: Sta (5.62%) | 1: Sta (34.69%) | 1: Sta (53.33%) | 1: Sta<br>(33.02%) | 1: Sta<br>(73.76%) |
| Vis | 1: Arl (35.36%)<br>2: Sta (15.39%)<br>3: Bur (14.29%)<br>9: Vis (1.07%) | 1: Jon (30.2%)<br>2: Bur (23.4%)<br>3: Lib (9.88%)<br>9: Vis (1.82%) | 1: Lib (27.67%)<br>2: Jon (20.94%)<br>3: Arl (15.93%)<br>6: Vis (7.76%) | 1: Vis<br>(83.98%) | 1: Vis<br>(71.37%) |
| Whe | 1: Boi (60.71%)<br>2: Bov (26.43%)<br>3: Whe (9.08%) | 1: Whe (99.99%) | 1: Whe (99.18%) | 1:Whe<br>(99.92%) | 1:Whe<br>(77.22) |

Table S7: Percentage match for misclassifications (Part 1 of 2) from geometric morphometrics using Feature Selection Random Forest. All cultivars with at least a single misclassification are included. The highest three percentages for each cultivar are included. If the correct classification is not in the top three then it is also included together with its rank. If the fruit was correctly classified only the correct classification posterior is included.

| Cultivar | Sample 1 | Sample 2 | Sample 3 | Sample 4 | Sample 5 |
|---|---|---|---|---|---|
| Ada | 1: Ada (51.6%) | 1: Ada (32.8%) | 1: Fuj (24.4%)<br>2: Ada (18.0%) | 1: Ada (19.8%) | 1: Ada (27.6%) |
| Bea | 1: Fuj (39.4%)<br>2: Bea (19.6%) | 1: Bea (53.2%) | 1: Bea (32.8%) | 1: Bea (43.6%) | 1: Bea (34.8%) |
| Boi | 1: Boi (31.8%) | 1: Boi (40.0%) | 1: Bov (32.8%)<br>2: Cat (12.8%)<br>3: Boi (12.6%) | 1: Cat (17.8%)<br>2: Bov (17.6%)<br>3: Boi (12.2%) | 1: Boi (40.6%) |
| Bov | 1: Whe (46.0%)<br>2: Lim (10.0%)<br>3: Fuj (8.4%)<br>10: Bov (2.4%) | 1: Boi (42.2%)<br>2: Bov (17.6%) | 1: Boi (41.8%)<br>2: Bov (18.4%) | 1: Boi (33.2%)<br>2: Bov (25.0%) | 1: Boi (26.0%)<br>2: Cat (19.6%)<br>3: Ada (17.6%)<br>4: Bov (15.6%) |
| Bur | 1: Bur (17.8%) | 1: Bur (30.4%) | 1: Cox (26.6%)<br>2: Bur (10.4%) | 1: Bur (37.2%) | 1: Bur (47.0%) |
| Cat | 1: Cat (58.4%) | 1: Ada (17.6%)<br>2: Cat (14.4%) | 1: Cat (53.8%) | 1: Cat (47.2%) | 1: Bov (28.2%)<br>2: Cat (20.6%) |
| Cha | 1: Cha (27.8%) | 1: Clo (37.4%)<br>2: Cha (28.4%) | 1: Cha (32.8%) | 1: Gra (35.0%)<br>2: Cha (22.6%) | 1: Cha (45.6%) |
| Cox | 1: Sta (21.2%)<br>2: Jon (15.8%)<br>3: Cox (10.6%) | 1: Gra (15.4%)<br>2: Cox (14.8%) | 1: Cox (22.2%) | 1: McI (21.4%)<br>2: Cox (14.6%) | 1: Cox (21.8%) |
| Flo | 1: Flo (34.6%) | 1: Flo (29.2%) | 1: Flo (49.8%) | 1: Sta (17.2%)<br>2: Bur (14.0%)<br>3: Vis (11.6%)<br>5: Flo (9.4%) | 1: Vis (26.2%)<br>2: Bra (20.4%)<br>3: Flo (8.2%) |
| Fuj | 1: Ada (23.2%)<br>2: Fuj (17.6%) | 1: Fuj (33.4%) | 1: Rhe (25.8%)<br>2: Fuj (21.8%) | 1: Fuj (41.8%) | 1: Fuj (43.4%) |
| Gol | 1: Gol (25.0%) | 1: Clo (24.2%)<br>2: Gra (17.8%)<br>3: Cha (15.8%)<br>5: Gol (8.4%) | 1: Sir (34.0%)<br>2: Gol (22.4%) | 1: Gol (31.8%) | 1: Gra (52.6%)<br>2: Gol (10.8%) |

Table S8: Percentage match for misclassifications (Part 2 of 2) from geometric morphometrics using Feature Selection Random Forest. All cultivars with at least a single misclassification are included. The highest three percentages for each cultivar are included. If the correct classification is not in the top three then it is also included together with its rank. If the fruit was correctly classified only the correct classification posterior is included.

| Cultivar | Sample 1 | Sample 2 | Sample 3 | Sample 4 | Sample 5 |
|---|---|---|---|---|---|
| Gra | 1: Jon (29.8%)<br>2: Arl (24.6%)<br>3: Lib (13.6%)<br>6: Gra (5.0%) | 1: Gra (32.4%) | 1: Gra (45.0%) | 1: Gra (36.4%) | 1: Clo (19.6%)<br>2: Gra (13.2%) |
| Jon | 1: Bur (19.6%)<br>2: Arl (17.8%)<br>3: Jon (14.6%) | 1: Lib (21.2%)<br>2: Arl (13.8%)<br>3: Jon (12.4%) | 1: Jon (22.2%) | 1: Arl (40.8%)<br>2: Jon (10.4%) | 1: Sta (17.0%)<br>2: Arl (13.0%)<br>3: Cox (10.5%)<br>4: Jon (9.8%) |
| Lib | 1: Lib (23.6%) | 1: Lib (23.8%) | 1: Lib (24.4%) | 1: McI (28.8%)<br>2: Flo (14.2%)<br>3: Lib (13.4%) | 1: Arl (19.8%)<br>2: Lib (18.4%) |
| Lim | 1: Lim (62.8%) | 1: Lim (65.0%) | 1: Lim (49.6%) | 1: Lim (62.6%) | 1: Kai (44.0%)<br>2: Lim (29.2%) |
| McI | 1: McI (36.8%) | 1: McI (34.2%) | 1: McI (17.8%) | 1: Jon (21.2%)<br>2: McI (15.6%) | 1: Bur (23.4%)<br>2: McI (16.0%) |
| Red | 1: Red (34.4%) | 1: Red (23.4%) | 1: Red (48.4%) | 1: Red (44.6%) | 1: Bea (33.0%)<br>2: Red (22.6%) |
| Rhe | 1: Rhe (61.6%) | 1: Rhe (38.2%) | 1: Rhe (50.2%) | 1: Rhe (39.0%) | 1: Fuj (28.0%)<br>2: Rhe (18.6%) |
| Sir | 1: Sir (28.4%) | 1: Cox (16.8%)<br>2: Sir (11.4%) | 1: Sir (37.0%) | 1: Gol (22.8%)<br>2: Sir (21.6%) | 1: Cox (39.8%)<br>2: Sir (13.0%) |
| Sta | 1: Sta (21.6%) | 1: Sta (30.6%) | 1: Sta (30.8%) | 1: Arl (19.2%)<br>2: Gra (17.2%)<br>3: Sta (5.6%) | 1: Flo (20.8%)<br>2: Sta (16.2%) |
| Vis | 1: Lib (33.4%)<br>2: McI (19.6%)<br>3: Flo (8.8%)<br>9: Vis (4.4%) | 1: Flo (24.4%)<br>2: Bra (19.0%)<br>3: Sta (10.2%)<br>7: Vis (2.8%) | 1: Vis (59.0%) | 1: Vis (44.6%) | 1: Vis (46.6%) |
| Whe | 1: Sta (11.2%)<br>2: Arl (9.2%)<br>3: Jon (7.6%)<br>16: Whe (0.2%) | 1: Whe (47.4%) | 1: Whe (38.6%) | 1: Whe (19.2%) | 1: Whe (33.8%) |
